# Microporous scaffolds drive assembly and maturation of progenitors into β-cell clusters

**DOI:** 10.1101/560979

**Authors:** Richard L. Youngblood, Joshua P. Sampson, Kimberly R. Lebioda, Graham Spicer, Lonnie D. Shea

## Abstract

Human pluripotent stem cells (hPSCs) represent a promising cell source for the development of β-cells for use in therapies for type 1 diabetes. Current culture approaches provide the signals to drive differentiation towards β-cells, with the cells spontaneously assembling into clusters. Herein, we adapted the current culture systems to cells seeded on microporous biomaterials, with the hypothesis that the pores can guide the assembly into β-cell clusters of defined size that can enhance maturation. The microporous scaffold culture allows hPSC-derived pancreatic progenitors to form clusters at a consistent size as cells undergo differentiation to immature β-cells. By modulating the scaffold pore sizes, we observed 250-425 µm pore size scaffolds significantly enhance insulin expression and key β-cell maturation markers compared to suspension cultures. Furthermore, when compared to suspension cultures, the scaffold culture showed increased insulin secretion in response to glucose stimulus indicating the development of functional β-cells. In addition, scaffolds facilitated cell-cell interactions enabled by the scaffold design and cell-mediated matrix deposition of extracellular matrix (ECM) proteins associated with the basement membrane of islet cells. We further investigated the influence of ECM on cell development by incorporating an ECM matrix on the scaffold prior to cell seeding; however, their presence did not further enhance maturation. These results suggest the microporous scaffold culture facilitates 3D cluster formation, supports cell-cell interactions, and provides a matrix similar to a basement membrane to drive *in vitro* hPSC-derived β-cell maturation and demonstrates the feasibility of these scaffolds as a biomanufacturing platform.

## 1. Introduction

Type I diabetes (T1D) is a chronic metabolic disorder characterized by autoimmune destruction of the pancreatic β-cells that results in the need for life-long insulin therapy. This disease represents 5–10% of the diagnosed cases of diabetes, corresponding to more than 1.5 million individuals in the United States[1]. Several secondary metabolic disorders can arise from this disease, as well retinopathy, neuropathy, nephropathy, stroke and heart failure[2, 3]. Although exogenous insulin injections have decreased mortality, hypoglycemic events and vascular complications persist[4–6]. Thus, recent research has turned to cell-based therapies focused on replacing lost insulin-producing cells. Enthusiasm in cell replacement therapies for diabetes was driven, in part, by the progress in allogeneic islet transplantation with the Edmonton protocol[7, 8]. However, the widespread application of islet transplantation has been tempered by the lack of availability of islets, the need for life-long immunosuppression, and the increasing reversal of insulin independence in as many as 50% of the recipients 5 years after the transplantation, although some β-cell function and hypoglycemia awareness seems to persist for longer than 5 years[9, 10].

The lack of available islets has led to the investigation of human pluripotent stem cells (hPSCs) as an unlimited source of functional β-cells. Multiple investigators have demonstrated the feasibility of differentiating hPSCs to immature β-cells *in vitro*, and transplanting these cells to support their maturation into glucose-responsive insulin-producing β-cells[11–17]. Initial findings from the Kieffer and Baetge/D’Amour groups demonstrated the production of pancreatic progenitors and, subsequently, immature β-cells *in vitro*, which could further differentiate following transplantation to normalize blood glucose levels after approximately 120 days[14, 18]. More recently, *in vitro* culture protocols have developed hPSC-derived β-cells that induce normoglycemia over shorter times after transplantation[12,14,19,20]. While numerous protocols have been established, the *in vitro* production of β-cells can result in a heterogenous population consisting of polyhormonal endocrine cells in addition to monohormonal β-cells. Moreover, the transplantation of these insulin-producing β-cells does not result in all recipients returning to normoglycemia[12, 13], indicating the need to more consistently and efficiently promote maturation of insulin-producing β-cells [17, 21]. Consistent and efficient methods for differentiation to monohormonal β-cells are required for clinical translation to treat T1D.[22]

The current stepwise hPSC differentiation approach aims to mimic a temporal control of organogenesis observed during embryonic development, and spatial control may represent an opportunity to enhance the efficiency and consistency of β-cell maturation. *In vivo* spatial control is achieved with cell-cell and cell-extracellular matrix (ECM) interactions. The ECM forms a three-dimensional (3D) environment and offers a niche for cell adhesion, migration, proliferation, and differentiation[23–25]. 3D environments have been shown to enhance differentiation and promote the assembly of functional tissues[12, 13]. Recent advances in three-dimensional cultures have generated tissues called organoids which possess several advantages including tissue structure and cellular organization similar to the native organ while possessing the specified cell types[26–32]. These same cell types cultured on plastic and allowed to simply self-assemble are not able to form the same complex tissue architectures that are permitted by 3D cultures.

Recently, microporous scaffolds have been used to provide a 3D environment to facilitate the assembly of hPSCs into multicellular structures, or organoids[33, 34]. Microporous scaffolds allow for both cell-cell and cell-matrix signaling that supports the self-organization of the cells into functional tissue structures[35]. This approach has proved groundbreaking for components of the lung, intestine, brain, liver, stomach, and the eye[26–32]. hPSC-derived β-cells have currently been obtained through culture as a monolayer of 2D clusters or as 3D aggregates on low attachment plates or in suspension cultures [12,14,17]. However, *in vivo* these cells naturally congregate into islets that are surrounded by a supportive extracellular matrix[36–38]. Mimicking the spatial cues in the pancreatic niche environment has the potential to augment the *in vitro* hPSC differentiation toward glucose-responsive, insulin-secreting cells.

Herein, we investigated microporous polymer scaffolds as an *in vitro* platform for the efficient differentiation of these progenitor cells to glucose-responsive insulin-producing β-cells[18, 39]. Scaffolds were formed from multiple synthetic polymeric materials, with the walls supporting the assembly of pancreatic progenitors into β-cell clusters. The pore size was investigated for the ability to form clusters of distinct sizes. The cell structures within the pores of the scaffold were analyzed by histology and gene expression. The influence of ECM-coated scaffolds on β-cell maturation was evaluated as well to investigate the role of the matrix in cellular assembly and differentiation. Finally, we assessed the maturation and function of these cells through glucose stimulated insulin secretion assays. These studies provide insight on cell-cell and cell-matrix interactions that influence the differentiation of hPSCs to β-cells on microporous scaffolds, which may ultimately provide a platform for biomanufacturing the cells as a therapy for T1D.

## 2. Materials and methods

### 2. 1 Microporous scaffold fabrication

Microporous scaffolds were fabricated as previously described[40, 41]. Two types of scaffolds were used poly(lactide-co-glycolide) (PLG) scaffolds and polyethylene glycol (PEG) scaffolds. Briefly, PLG microporous scaffolds were fabricated by compression molding PLG microspheres (75:25 mole ratio D,L-lactide to glycolide) and micron-sized salt crystals in a 1:30 ratio of PLG microspheres to salt. The mixture was humidified in an incubator for 7 min and then thoroughly mixed again. Scaffolds were compression molded with 77.5 mg of polymer–salt mixture into cylinders 5 mm in diameter by 2 mm in height using a 5 mm KBr die (International Crystal Laboratories, Garfield, NJ) at 1500 psi for 45s. Molded constructs were gas foamed in 800 psi carbon dioxide for 16 h in a pressure vessel. The vessel was depressurized at a controlled rate for 30 min. On the day of cell seeding, scaffolds were leached in water for 1.5 h, changing the water once after 1 h. Scaffolds were sterilized by submersion in 70% ethanol for 30 seconds and multiple rinses with phosphate buffer solution.

For the PEG hydrogels, PEG-maleimide (4-arm, molecular weight 20kDA, 20% wt/wt) polymer was dissolved in a HEPES buffer solution, mixed with NaCl crystals and a photoinitiator (Irgacure-2959) then cast into a PDMS mold (diameter: 5mm, thickness 2mm). The solution is irradiated with UV light to photo-crosslink the PEG-maleimide and then washed to remove the sodium chloride and unreacted photoinitiator. The pore size of the scaffolds can be readily controlled through the dimensions of the porogen, and we propose to investigate pore sizes of 63 to 108 µm, 108-225 µm, 225 to 450 µm, and 500 to 600 µm.

The differentiation of pluripotent hPSCs (Stage 0) to pancreatic progenitor cells (Stage IV) were performed as described[14]. Cells were seeded on scaffolds at concentrations ranging from 12.5-125 million cells/cm^3^ and distributed across both faces of the scaffold. The differentiation towards immature β-cells (Stages V-VI) required 1-2 weeks based on established protocols[12,14,42] The impact of microporous scaffolds on the differentiation was compared to the established suspension culture method.

### 2.2 Protein adsorption to scaffolds

For coating scaffolds with ECM proteins, scaffolds were fabricated and disinfected in 70% ethanol and dried again before being placed into individual wells of a 24-well tissue culture dish. Proteins were then coated per manufacturer’s recommendations and to be consistent with our previous report that demonstrated enhancement in islet function following transplantation on ECM protein-modified scaffolds[43, 44]. Collagen IV (1mg/mL, Sigma), laminin-332 (1mg/mL, formerly termed laminin-5 and hereafter referred to as “laminin”, Sigma), Matrigel (Corning, Cat#: 354277) or PBS were added to the scaffold. The scaffolds were then incubated at 37°C for 1 h, followed by a second coating of the same component to each scaffold. Scaffolds were incubated with 95% humidity at 37°C overnight to facilitate protein adsorption to the scaffold surface.

### 2.3 Cell culture

We used a six-stage differentiation strategy to differentiate hPSCs in to pancreatic progenitors and further into β-cell clusters within the scaffold. For the suspension culture control, undifferentiated hPSCs were initially seeded at 1.0 million cells/1mL in ultra-low attachment 6 well plates (Corning, VWR) placed on a 9-Position stir plate (Chemglass) set at rotation rate of 95 rpm in a 37°C incubator, 5% CO2, and 100% humidity.

### 2.4 Cell viability

The viability of the cells was qualitatively assessed using Live/Dead® viability/cytotoxicity kit (Life Technologies), visualizing viable and dead cells by staining with the fluorescent dyes calcein and ethidium homodimer-1 (EthD-1), respectively. Cells cultured on microporous scaffolds were stained with 4 µM EthD-1 and 2 µM Calcein AM for 30 min before the viability was assessed using fluorescence microscopy.

### 2.5 qRT-PCR Analysis

For gene expression analysis, cell containing scaffolds were mechanically homogenized in a tube containing Trizol® reagent (Life Technologies), and RNA was isolated according to the manufacturer’s instructions. RNA concentration was determined using a NanoDrop spectrophotometer. The iScript™ Reverse Transcription Supermix was used to transcribe RNA into cDNA. SYBR Green dye was used to detect fluorescence. The amplification profile was assessed using a LightCycler® 480 (Roche, Germany). Gene expression was quantified using the ΔΔCt method and fold change was calculated using the formula 2^-ΔΔCt^. Values for the genes of interest were normalized to the housekeeping gene (GAPDH) followed by normalization to marker expression in pluripotent hPSCs. Primers used for qPCR analysis are listed in **Table 1**.

**Table 1.**
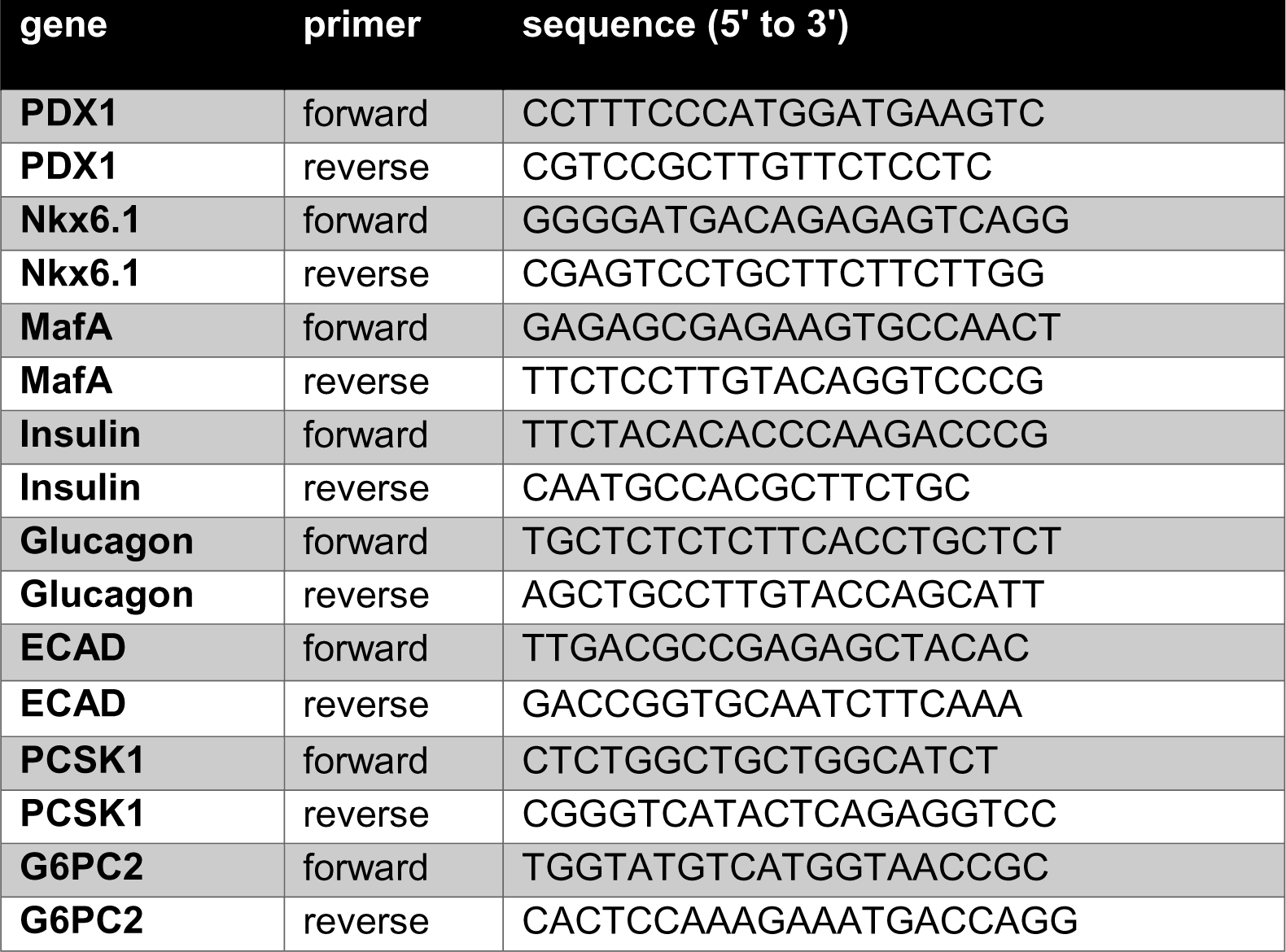
Primers of Pancreatic Differentiation Markers Used for qRT-PCR.

### 2.6 Immunostaining

Immunostaining of *in vitro* cell differentiation was performed at stage 6 (immature beta cells). Cells differentiated in suspension cultures were fixed using 4% paraformaldehyde (Electron Microscopy Sciences; Hatfield, PA, United States), permeabilized using 0.5% Triton in tris-buffered saline, blocked using normal donkey serum, and stained for cell-cell and cell-matrix interaction markers ECAD, Collagen IV, Laminin, Fibronectin as well as insulin and cell nuclei were counterstained with DAPI. Scaffold cultures were cryopreserved in isopentane cooled on dry ice and then embedded within OCT embedding medium and cryosectioned. Digital images were acquired with a MicroFire digital camera (Optronics, Goleta, CA) connected to an Olympus BX-41 fluorescence microscope (Olympus, Center Valley, PA, United States). Image quantification was conducted using MATLAB software using an object-based colocalization analysis. DAPI+ cells were identified out of the total area of the sectioned tissue and quantified by applying Otsu’s thresholding method, the watershed transform, and individual cluster thresholding. Then, each cell’s colocalization with immunofluorescent markers was quantified. For confocal imaging, whole tissue samples were fixed in 4% paraformaldehyde then stained with primary and secondary antibodies as described above. The labeled samples were then cleared in Murray’s clear solution for optical clearing for 45 min before being imaged via confocal microscopy (Nikon A1Si laser scanning confocal microscope, Nikon Instruments Inc, Tokyo, Japan).

### 2.7 Static Glucose-Stimulated Insulin Secretion Assay

We investigated the function of hPSC-derived β-cells *in vitro* under all scaffold conditions by a glucose stimulated insulin secretion assay. Immature β-cells derived from hPSCs in vitro have a capacity for glucose-stimulated insulin secretion (GSIS), though the magnitude of response may not be the same as fully mature functional β-cells. To analyze GSIS, scaffold cultures were first exposed to basal levels of glucose (2.8 mM), which was dissolved in Krebs-Ringer-Bicarbonate (KRB) buffer, and incubated for an hour. Samples of the buffer were collected to measure insulin levels using ELISA. The scaffolds were then washed in fresh basal level glucose and next exposed to a high-level glucose concentration (28 mM) for an hour. Samples of the buffer were collected for ELISA as outlined above. The quantification of insulin secretion at the various glucose exposures represent the GSIS.

### 2.8 Statistics

All statistical analyses were conducted using Prism graphing and data analysis software (GraphPad Software, Inc., La Jolla, CA, United States). Values were reported as the mean ± SEM.

## 3. Results

### 3. 1 Cluster formation of hPSC-derived pancreatic progenitors within scaffold pores

Two commonly used biomaterials, PLG and PEG, were investigated for the *in vitro* scaffold culture of pancreatic progenitors to β-cells. These two materials provided a similar microporous structure (**Fig 1A**) with distinct material properties that can distinguish the role of structure relative to the role of the material. We first evaluated cluster formation within the microporous scaffold cultures (**Fig 1B**) compared to the traditional suspension culture (**Fig 1C**). hPSC-derived pancreatic progenitors dissociated into single cells were initially seeded onto microporous scaffolds at a density of 12.5 million cells/cm^3^ for culture. Through confocal microscopy, we found that this seeding density was not sufficient for cluster formation to occur as cells were localized to the surface of the pores (**Fig 1D**). Increasing the seeding density 10-fold to 125 million cells/cm^3^ resulted in the cells assembling into 3D clusters within one day after seeding (**Fig 1E**), which resembles the self-organization that occurs in suspension cultures. These observations indicated that the assembly of cells into clusters could be supported within the micropores, yet was dependent on the cell density. This high cell seeding density resulted in a uniform distribution of cells and clusters across both scaffold surfaces. At densities greater than 125 million cells/cm^3^, cells began to clump on the surface. Cell viability was consistently high (>90%) throughout the 14-day experiment, and similar for all scaffold conditions (**Fig 1F-K**). These results demonstrated that the pores could facilitate cluster formation.

**FIG 1:**
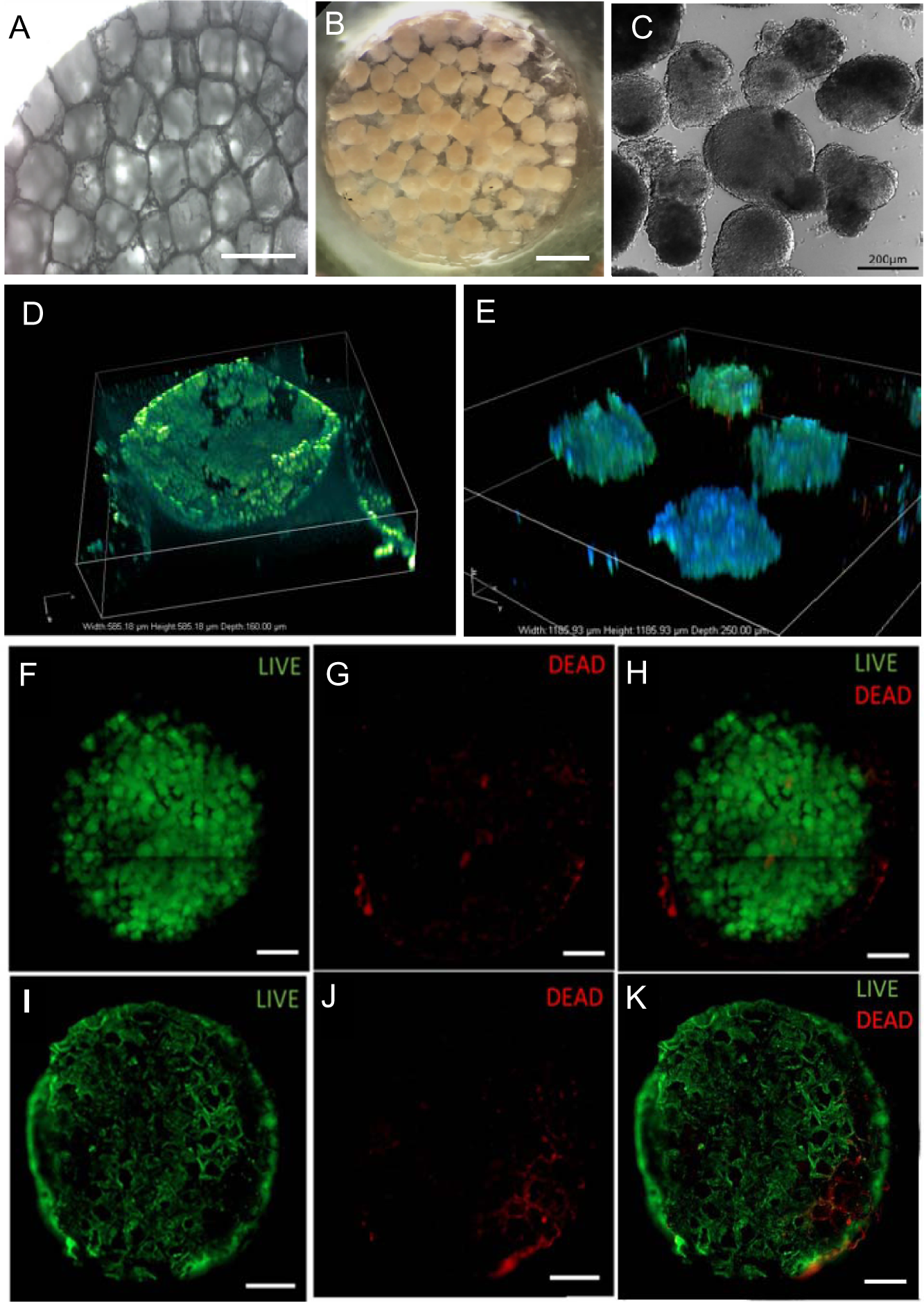
*In vitro* culturing of hPSC-derived pancreatic progenitors on microporous scaffolds. SEM image of empty porous PLG scaffold with 250-425 µm pores (A). Microscope images of PEG scaffold culture (B) and suspension culture (C). Confocal imaging shows cell localization within a scaffold pore at 12.5×10^6^ cells/cm^3^ (D) and multiple scaffold pores at 125×10^6^ cells/cm^3^ (E). β-cell progenitors were stained 2wks after seeding onto a PEG scaffold (F-H) and PLG scaffold (I-K) and examined using a live/dead assay. (Scale bar is 1 mm)

### 3. 2 Maturation of β-cell clusters within scaffolds

We next investigated the feasibility of generating β-cell clusters in microporous scaffolds by measuring the expression of β-cell marker genes, which were compared with cells generated in suspension culture and human islets. Using real-time quantitative PCR (qRT-PCR) analysis, we found cells cultured within both PEG and PLG scaffolds had significant increases in expression of endocrine hormone marker genes (*INS and GLUC*) relative to Stage IV pancreatic progenitors. Additionally, β-cell maturation markers (*MafA, G6PC2, and PCSK1*) had expression levels on scaffold cultures that were comparable to suspension culture controls. We also used the tunable design of the scaffold to assess how varying the pore size would influence maturation. A correlation between scaffold pore sizes and the expression of key β-cell markers was observed, with larger pore sizes resulting in better maturation. For PEG scaffolds, pore sizes in the range of 250-425 µm and 500-600 µm had expression of pancreatic transcription factors (*PDX1* and *Nkx6.1*) (**Fig 2A**) that was comparable to human islets. For scaffolds with pore sizes smaller than 250 µm, the expression of *Nkx6.1*, a marker for mature β-cells, was lower than human islets and suggested the cells have not fully matured. Additionally, relative to pancreatic progenitors, the expression of insulin only had a significant increase on scaffolds with pore sizes of 250-425 µm and 500-600 µm. This trend between scaffold pore size and cell development was observed in the expression of β-cell maturation markers as well. Scaffolds with a pore size of 250-425 µm resulted in the highest expression of the insulin gene transcription factor, *MafA,* with the 500-600 µm pore size scaffolds exhibiting the second highest expression. The maturation marker proprotein convertase 1 (*PCSK1*) is one of the key enzymes associated with insulin processing and showed expression levels to be higher in scaffold cultures with 250-425 µm pore sizes compared to the other scaffold pore sizes as well as suspension clusters.

**FIG 2:**
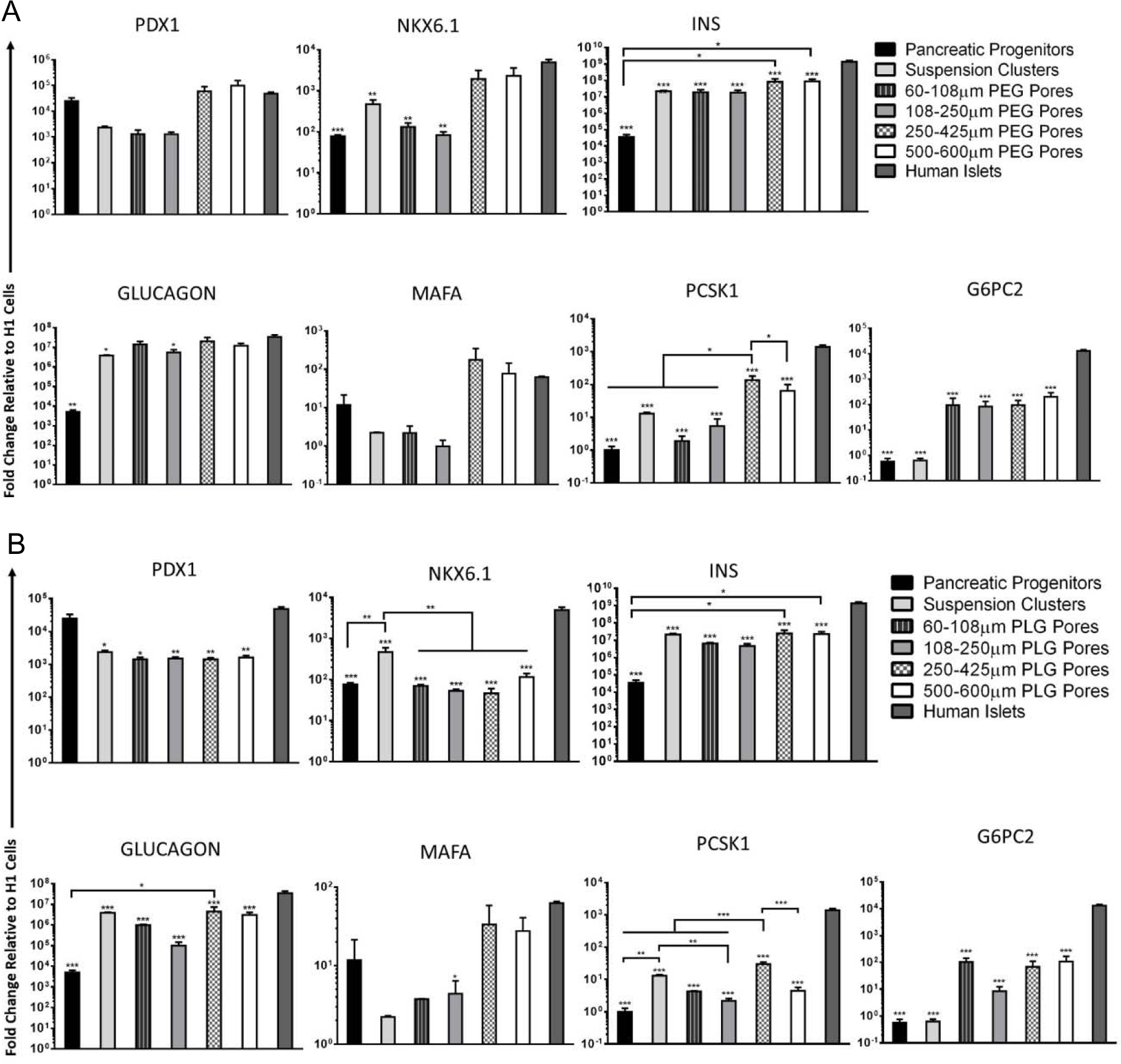
Augmenting Scaffold Pore Sizes Enhances β*-cell* Maturation. Gene expression profile of hPSC-derived cells as Stg 4 pancreatic progenitors and Stg 6 immature β-cells cultured on (A) PEG and (B) PLG microporous scaffolds with varying pore sizes. (n = 4 biological replicates for all genes). *P < 0.05, **P≤0.01, ***P≤0.001 for each condition versus human islets (one-way ANOVA with Dunnett test for multiple comparisons). Error bars represent the standard error of mean (SEM).

hPSC-derived pancreatic progenitors cultured on PLG scaffolds similarly demonstrated the development of maturing β-cells with maturation showing a correlation with pore size (**Fig 2B**). Insulin expression for PLG scaffold cultures with 250-425 µm pore sizes was significantly increased relative to the pancreatic progenitors. The expression of pancreatic β-cell transcription factor *PDX1* in all four PLG scaffold conditions were lower than human islets but comparable to suspension clusters. On the other hand, *Nkx6.1* expression was significantly lower in PLG scaffolds when compared to the suspension culture. While this suggests maturation could be improved, PLG scaffolds with 250-425 µm pore sizes did show increased expression in the key maturation marker, *PCSK1*, relative to pancreatic progenitors. Overall, cells cultured in PEG and PLG microporous scaffolds generally showed increased expression levels of β-cell maturation markers. While the maturation was still not comparable to human islets, this deficit relative to islets is to be expected as *in vivo* transplantation is necessary to reach full maturation. This analysis revealed that the differentiation of pancreatic progenitors to β-cells cultured in the PLG and PEG microporous scaffold was, at a minimum, comparable and improved under some conditions to the traditional suspension culture. Our observations identified a relationship between scaffold pore size and β-cell maturation with 250-425 µm pore size showing more significant improvements in β cell development than in suspension culture. Thus, the following studies focused on scaffolds with a pore size of 250-425 µm.

### 3.3 Cell-cell communication in microporous scaffold cultures during β-cell maturation

Next, we investigated cell-cell interactions within the scaffold culture that drive maturation in the pancreatic niche environment. The cell surface adhesion protein epithelial cadherin (ECAD) plays a critical role in the development of islets and intra-islet communication and is implicated in efficient insulin secretion from β-cells[45, 46]. Thus, we assessed the presence of ECAD through qRT-PCR and immunostaining in both scaffold cultures and suspension clusters. ECAD gene expression levels in hPSC-derived immature β-cells cultured in microporous scaffolds had a closer resemblance to the expression in human islets than suspension clusters (**Fig 3A**). PEG scaffold cultures had a higher expression level compared to suspension clusters. However, compared to human islets, both PEG (2.84±0.96 vs 6.70±1.22, n=3, *P*<0.05) and suspension culture (0.57±0.03 vs 6.70±1.2, n=3, *P*<0.01) conditions showed lower ECAD gene expression. The PLG scaffold culture (3.05±1.21, n=4) had expression that increased relative to suspension culture and was comparable to human islets. The promotion of cell-cell interactions in scaffolds was confirmed with immunostaining. E-cadherin protein expression was shown to be localized to specific regions in the suspension clusters (**Fig 3B**) whereas, in scaffold cultures, expression was distributed throughout the clusters (**Fig 3C**). Imaging analysis of DAPI^+^ cells colocalized with the cell-cell adhesion marker, ECAD, per the total area revealed PLG scaffold cultures increased protein expression of ECAD compared to suspension (42.8% ± 5.1 vs 11.2% ± 2.9 of total cell population, n=4; P<0.01). Our data suggests that microporous scaffold cultures, particularly PLG, promote cell-cell interactions that can play a role in driving β-cell maturation.

**FIG 3:**
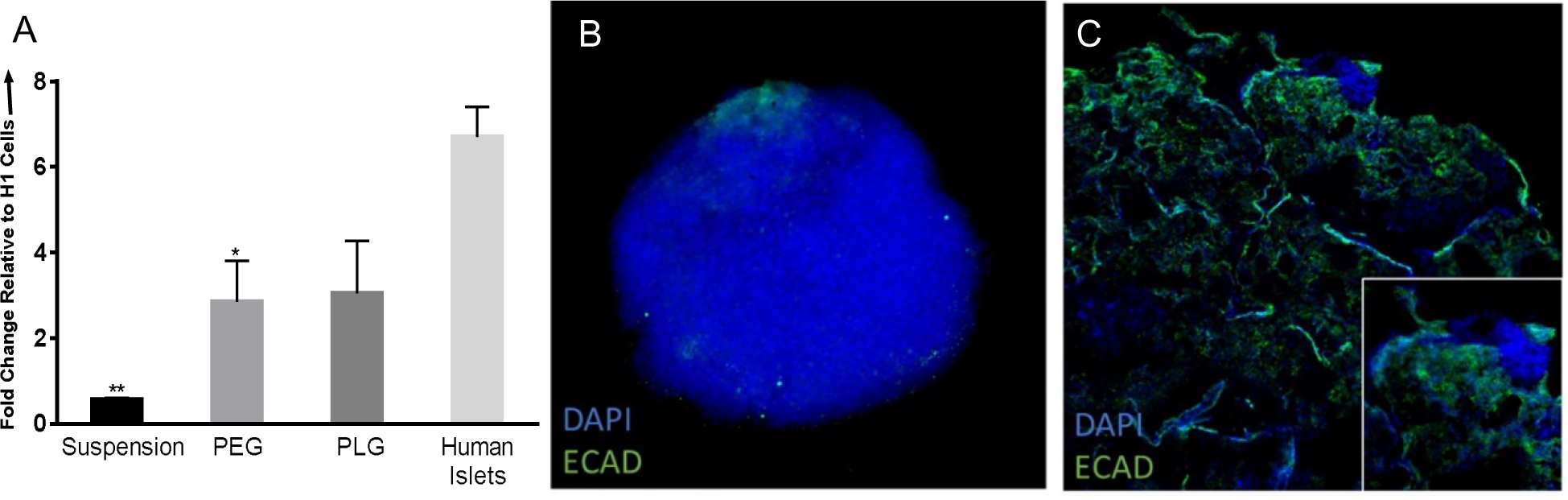
Scaffold Culture Influences E-cadherin Interactions In β-cell Clusters. ECAD gene expression levels of hPSC-derived immature β-cells cultured in suspension or on PEG and PLG microporous scaffolds (n = 3-4 biological replicates). *P < 0.05, **P<0.01 compared to human islets (one-way ANOVA with Dunnett test for multiple comparisons). Error bars represent the SEM. (A) Immunofluorescent staining of suspension cluster (B) and PLG scaffold culture (C) for ECAD (green) and DAPI (blue) with an inset of a scaffold pore.

### 3.4 hPSC-derived β-cell glucose-responsive in vitro function within microporous scaffolds

The function of these β-cell clusters in scaffolds was next examined by their ability to secrete insulin in a glucose-responsive manner. Scaffold cultures and suspension clusters at the end of the six-stage differentiation were exposed to 2.8 mM and 28 mM glucose solutions, respectively. We observed a sharp increase in insulin secretion from the scaffold cultures when exposed to a high concentration of glucose. hPSC-derived β-cells cultured on the PLG scaffold had the highest insulin secretion index, with a threefold increase compared to the suspension culture control (1.34 ± 0.20 vs 0.43 ± 0.06, n=3, *P*<0.01) (**Fig 4**). PEG scaffold cultures also showed higher insulin secretion compared to the suspension control (1.11 ± 0.09, n=3, *P*<0.05) yet lower insulin secretion than PLG. These results indicate that hPSC-derived β-cells cultured on microporous scaffold cultures are capable of glucose-stimulated insulin secretion, which represents functional maturation. Since the PLG scaffold demonstrated a higher degree of function compared to PEG, subsequent studies employed this material for further investigation of the matrix environment supporting β-cell maturation.

**FIG 4:**
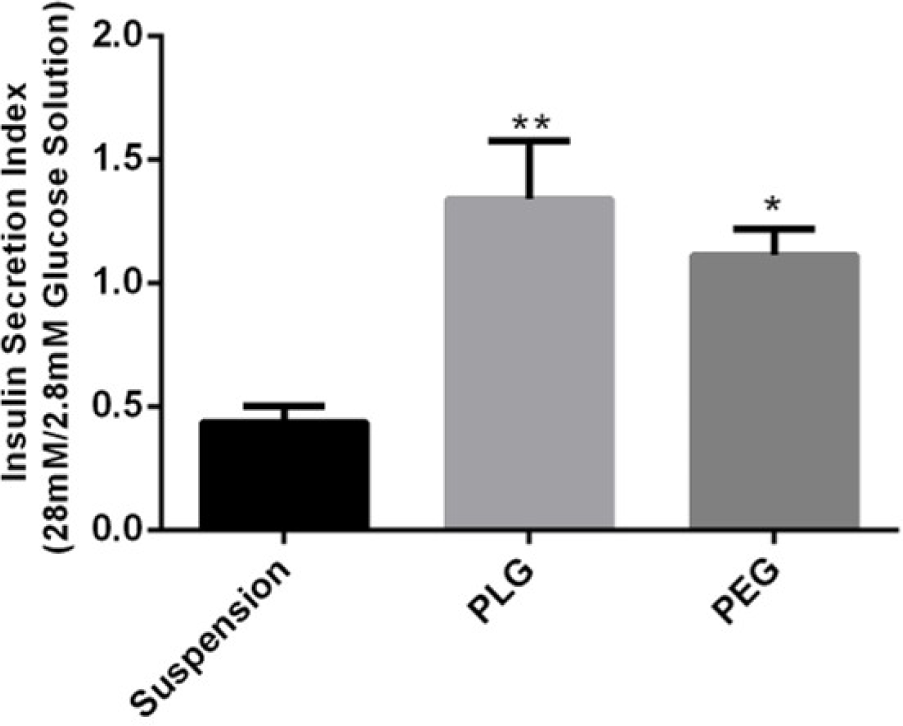
Efficient Generation of Glucose-Responsive Beta Cells from Human Pancreatic Progenitors Differentiated on Microporous Scaffold Cultures. Human insulin secretion from PLG and PEG scaffold cultures and suspension clusters in response to low and high glucose concentrations under static conditions (scaffold cultures: n = 3 biological replicates, suspension: n = 4 biological replicates). *P < 0.05, **P<0.01 compared to suspension. Error bars represent the SEM.

### 3.5 ECM deposition by hPSC derived β-cell clusters

We subsequently assessed the ECM within the cultured cells, as ECM basement membrane proteins are a critical component of the pancreatic environment supporting islets. Since cells cultured on PLG scaffolds showed signs of islet-like development and function, we used immunofluorescence analysis to investigate if the cells were establishing a matrix similar to the pancreas within the scaffold. Our results showed ECM proteins, collagen IV, laminin and fibronectin, were present in both suspension clusters (**Fig 5A-C**) and in PLG scaffold cultures (**Fig 5D-F**). Imaging analysis confirmed that the percentage of DAPI^+^ cells localized to ECM protein expression in the PLG scaffold cultures were comparable to suspension (collagen IV: 43.5%±3.5 vs 42.5%±11.8; n=4; laminin: 42.5%±6.2 vs 50.9±2.3, n=3; fibronectin: 41.1%±5.1 vs 50.7±4.3, n=3) (**Fig 5G**). The distribution of ECM protein across the cell cluster were analyzed using a statistical autocorrelation function, which determines the correlation length (L_c_) (**Fig 5H)**. For our purposes, the L_c_ quantifies the heterogeneity of image texture to show if the fluorescently-labeled protein was concentrated in large clumps or distributed across smaller regions. The L_c_ of PLG scaffold cultures was relatively low, indicating that ECM proteins were clustered in small, well distributed regions, while the suspension cultures had a relatively high L_c_ suggesting the ECM proteins were clumped into larger structures (18.2 µm ± 3.2 vs 68.3 µm ± 10.0, n=3, P<0.0001). Thus, the scaffolds had a more homogeneous distribution of ECM proteins throughout the β-cell clusters compared to suspension culture.

**FIG 5:**
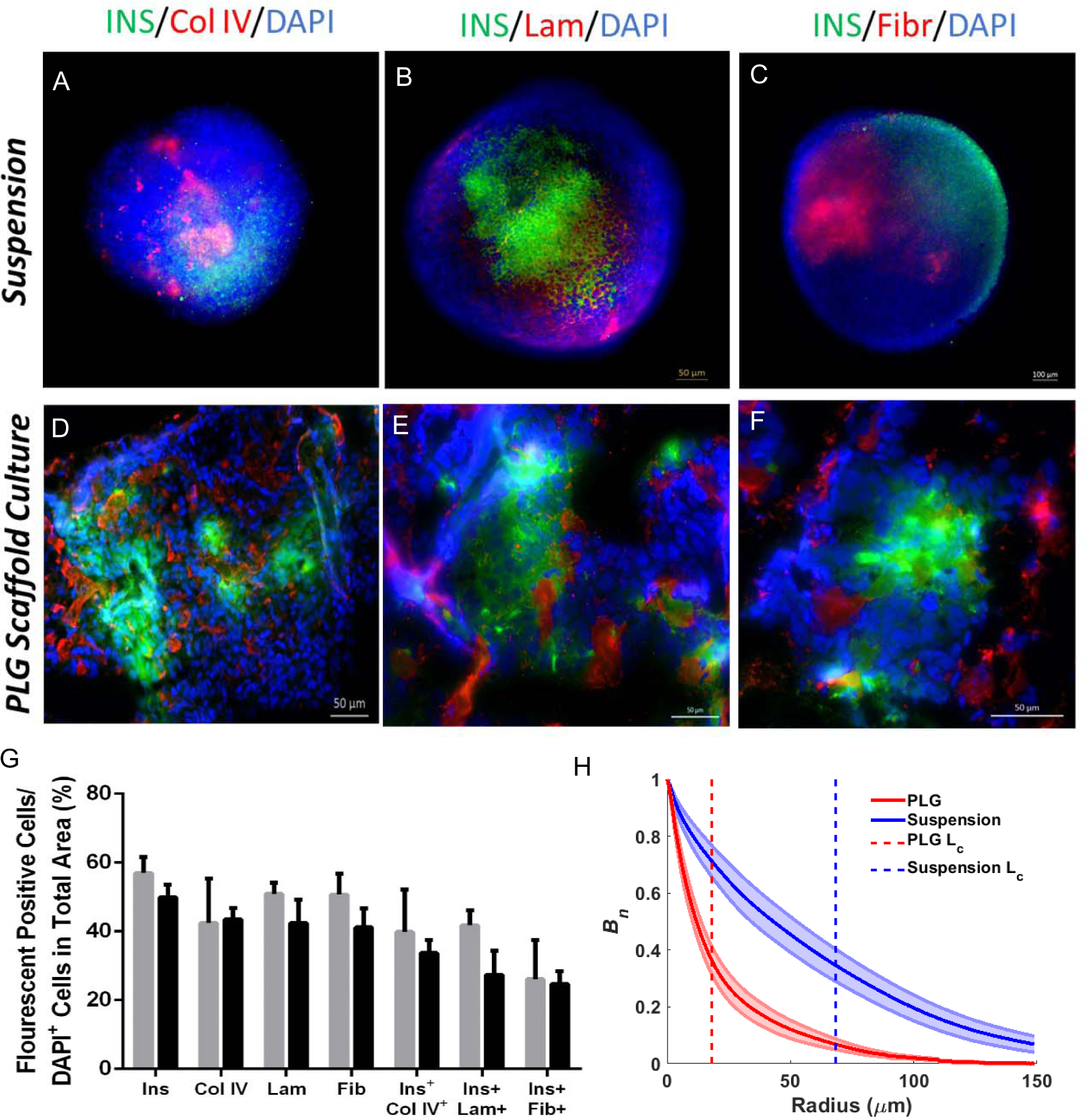
Scaffold Cultures Permit hPSC-derived β-cell-Secreted ECM Deposition. Representative immunofluorescent staining of suspension cluster (A-C) and PLG scaffold culture (D-F) for insulin (green), ECM protein (collagen IV, laminin, or fibronectin) (red) and DAPI (blue). Scale bar is 100 µm. (G) Immuno-histological analysis of β-cells in suspension (gray) and PLG microporous scaffolds (black) for the percentage of cells expressing insulin, ECM proteins and insulin^+^ cells colocalized with ECM proteins (n = 3-4 biological replicates) with error bars representing SEM. (H) Autocorrelation functions are plotted for immunoflourescent images of DAPI^+^ cells expressing ECM protein on either PLG scaffolds (red) or suspension (blue) cultures. A dotted line for each function marks the correlation length L_c_, the distance (in microns) at which the image autocorrelation function decays to 1/e of its initial value at r=0. SEM is plotted as regions around each autocorrelation function (n=3 biological replicates).

### 3.6 β-cell maturation on ECM-modified microporous scaffolds

The presence of ECM protein deposition on the naked scaffold motivated studies in which ECM proteins commonly found in the pancreas were deposited on the scaffold prior to cell seeding as a means to further enhance maturation. Using qRT-PCR analysis, we investigated pancreatic progenitor maturation to β-cells on PLG scaffolds coated with either Collagen IV, Laminin or Matrigel. Naked microporous scaffolds and scaffolds coated with ECM proteins showed comparable levels of expression for endocrine transcription factors (*PDX1 and Nkx6.1)* (**Fig 6A**). However, only naked scaffolds and Matrigel coated scaffolds exhibited an increase in insulin expression relative to pancreatic progenitors. Evaluating cell maturation on the scaffolds also showed that, relative to pancreatic progenitors, only the naked scaffold cultures enhanced the expression of *PSCK1*. An analysis of the expression of pancreatic-related ECM genes demonstrated that the expression of *COL4A1* gene, coding for collagen type IV, in naked scaffolds as well as scaffolds coated with Collagen IV and Laminin was comparable to human islets. However, a decrease of *COL4A1* gene expression was observed in suspension cultures and scaffolds coated with Matrigel (**Fig 6B**). Laminin production, indicated by *LamA5* gene, was observed to be comparable to human islets across all culture conditions. Finally, we assessed maturation through glucose-responsive function and, interestingly, observed scaffold cultures pre-coated with ECM proteins did not show a significant improvement in β-cell function compared to naked scaffolds (0.63± 0.03 for PLG with Matrigel, 1.15 ± 0.15 for PLG with Laminin, 1.02 ± 0.17 for PLG with Collagen IV compared to 1.34 ± 0.20 for PLG, n=3) (**Fig 6C**). These findings indicate initial introduction of ECM does not substantially improve maturation at the end of the scaffold culture, which could be potentially due to deposition of matrix proteins by cells throughout the culture.

**FIG 6:**
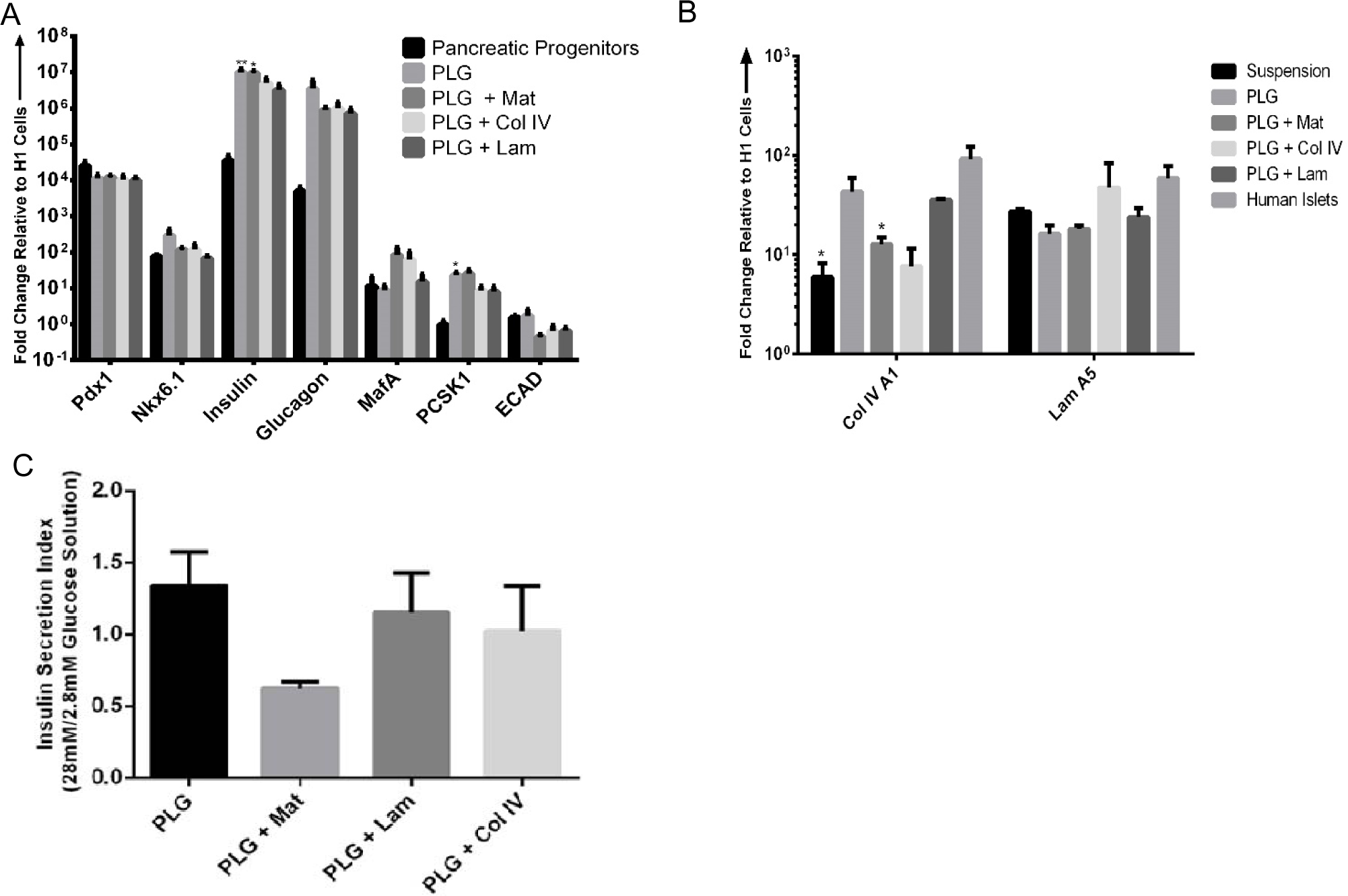
*In Vitro* Extracellular Matrix Development in hPSC-derived β-cell Clusters. Gene expression profile of Stg6 immature β-cells cultured on ECM-coated microporous PLG scaffolds. (n = 4 biological replicates for all genes). *P < 0.05, **P<0.005 versus pancreatic progenitors. (A) Gene expression for ColIVA1 and LamA5 of Stg6 immature β-cells. *P < 0.05 compared to human islets (Student t-test comparisons) (B) Human insulin secretion in response to low and high glucose concentrations from PLG scaffold cultures coated with either Matrigel (Mat), Laminin (Lam) or Collagen IV (Col IV) under static conditions (n = 3 biological replicates,). *P < 0.05 compared to PLG. (C) Error bars represent the SEM.

## 4. Discussion

Microporous polymer scaffolds formed from synthetic materials can support the differentiation of hPSC-derived pancreatic progenitors toward immature β-cells *in vitro.* We chose scaffolds made from synthetic materials as they have been widely used for islet transplantation[43,44,47], and are generally easy to manufacture. This microporous architecture provides the opportunity to control and support the 3D organization of cells into β-cell clusters. hPSC-derived β-cells cultured on the scaffold showed significantly increased gene expression of pancreatic endocrine hormones, INSULIN and GLUCAGON, relative to pancreatic progenitors. Furthermore, the expression of β-cell maturation markers (*MAFA, PCSK1*, and *G6PC2*) was increased on the scaffold compared to the suspension clusters. β-cell maturation was further investigated through glucose-responsive functional tests that demonstrated cells cultured on the scaffold had higher insulin secretion than suspension clusters. The high porosity and interconnected pores of the scaffold may allow clusters to communicate through paracrine factors similar to the pancreas tissue[48, 49]. Despite their similar biomaterial design, the microporous scaffolds also exhibited a few differences in how they influenced cell differentiation. PEG scaffold cultures showed a more significant increase in the expression of β-cell maturation markers than PLG scaffolds. On the other hand, ECAD expression suggested that the PLG scaffolds supported cell-cell interactions comparable to islets, which could have played a role in the higher insulin secretion observed in PLG scaffolds versus PEG scaffold cultures. Despite the difference in materials, the architecture of the material supported the formation of β-cell clusters, suggesting a role of the material structure in enhancing maturation.

The microporous scaffold provides an environment conducive to controlling the size of the structures and promoting cell-cell connectivity that is essential for maturation. The size of transplanted islets has been previously reported to impact insulin secretion and viability[50–52]. Small islet clusters can exhibit low amounts of insulin secretion, which has been attributed to limited cell-cell contact, while excessively large clusters are considered to have limitations from nutrient availability[53]. Based on these results with islets, the influence of pore size, which would determine the hPSC-derived β-cell cluster size, was investigated. Protocols generally rely on hPSCs to spontaneously cluster in suspension resulting in the end-stage clusters varying in size, though efforts have started to focus on establishing a mechanism for controlling the size of the transplanted clusters due to the influence on long-term viability and the secretion of sufficient insulin[53]. Physical manipulation and shear have largely been employed to provide control. The pores of the scaffold can provide a direct control on cluster size. Our results suggest that clusters forming within 250-425 µm pores maximally support maturation toward β-cells. The pore size, in combination with seeding density, are also critical for promoting cell-cell interactions[54]. At low seeding densities, the cells were observed to primarily attach to the walls of the pores. However, increasing the cell density increasingly favored cell-cell interactions within the pore. Expression of E-cadherin was increased within scaffold culture relative to suspension culture, and E-cadherin staining was observed primarily between cells within the pores and not at the material surface. E-cadherin is a key player in maturation as studies have shown that E-cadherin immune-neutralization reduces both basal and glucose-stimulated insulin secretion[45]. Collectively, the microporous scaffold can be employed to control the formation of clusters, and to favor cell-cell interactions that are influential in maturation.

In addition to cell-cell interactions, interactions between stem cells and the extracellular matrix can induce lineage-specific differentiation and support the function of differentiated cells by providing a composite set of chemical and structural signals[55]. Herein, we report that differentiating cells deposit ECM proteins within the scaffold, with the composition resembling that found in the basement membrane around islets. Furthermore, relative to suspension culture, we found ECM proteins were more homogeneously distributed for the scaffold culture, with ECM proteins available for interaction with β-cells throughout the cluster. The suspension clusters had ECM deposited in relatively large clumps as indicated by the increased L_c_. Matrix deposition by the cells is likely a key step in maturation, as key integrins change over developmental stages[56]. This deposition of matrix initiates the formation of a niche, which is normally present in islets and influences maturation and function. Attachment of cells to ECM may also benefit β-cells by maintaining tissue architecture and preserving specific intercellular relationships within the pores. Interestingly, scaffolds coated with either collagen IV, laminin or Matrigel showed comparable maturation to naked scaffolds that relied on cell-secreted ECM, which contained collagen IV, laminin and fibronectin. The analysis of glucose-responsive insulin secretion on ECM coated scaffolds and naked scaffolds showed β-cell function was similar across all conditions as well.

## 5. Conclusion

This research demonstrates that scaffolds may function as a platform for manufacturing β-cells for transplantation. Suspension cultures have been used for aggregated hPSC-derived β-cell production, without the need for feeders and using procedures that are scalable to generate sufficient cells. However, hydrodynamic shear force-related cellular damage presents a challenge for suspension cultures generally[57]. Furthermore, the increasing culture volumes can influence the size of cell aggregates, which has previously been linked to apoptosis-related cell loss, cellular differentiation, and heterogeneity[58]. Culture on scaffolds represents an alternative manufacturing platform that has been employed for tissue engineered products[59]. The pore size can be employed to control the size of aggregates, and similarly the culture on scaffolds may also decrease the effects of hydrodynamic shear on clusters[60]. The scaffold design can be tuned to control cluster size, with ECM deposition on the supportive matrix. Similar to organoid development in three-dimensional cultures[26–32], the cells establish a functional niche during *in vitro* culture that may enhance engraftment *in vivo*. Furthermore, the scaffolds are formed from materials that have been used *in vivo*, and thus the cell-material construct could be directly transplanted, which has the added advantage of maintaining the niche that has developed within the scaffold. Collectively, scaffolds have the potential to serve as a component of the manufacturing of β-cells for transplantation.

